# Synonymous sites in SARS-CoV-2 genes display trends affecting translational efficiency

**DOI:** 10.1101/2020.05.30.125740

**Authors:** Qingsheng Huang, Huan Gao, Lingling Zheng, Xiujuan Chen, Shuai Huang, Huiying Liang

## Abstract

A novel coronavirus, SARS-CoV-2, has caused a pandemic of COVID-19. The evolutionary trend of the virus genome may have implications for infection control policy but remains obscure. We introduce an estimation of fold change of translational efficiency based on synonymous variant sites to characterize the adaptation of the virus to hosts. The increased translational efficiency of the M and N genes suggests that the population of SARS-CoV-2 benefits from mutations toward favored codons, while the ORF1ab gene has slightly decreased the translational efficiency. In the coding region of the ORF1ab gene upstream of the −1 frameshift site, the decreasing of the translational efficiency has been weakening parallel to the growth of the epidemic, indicating inhibition of synthesis of RNA-dependent RNA polymerase and promotion of replication of the genome. Such an evolutionary trend suggests that multiple infections increased virulence in the absence of social distancing.

## INTRODUCTION

Severe acute respiratory syndrome coronavirus 2 (SARS-CoV-2), a previously unknown coronavirus, has caused the pandemic of coronavirus disease 2019 (COVID-19) since December 2019(*1–4*). Coronaviruses are positive-sense single-stranded RNA viruses with envelopes. The reference genome of SARS-CoV-2 (NCBI accession NC_045512) has 29,903 nucleotides (nt) containing ORF1ab (also known as NSP, non-structural protein), S (surface or spike), E (envelope), M (membrane or matrix), N (nucleocapsid), and other seven putative open reading frames (ORFs)(*2, 3*). Comparisons between genomes of SARS-CoV-2 and related viruses have revealed many important insights about the origin of SARS-CoV-2(*5–8*), but there are few studies on how the genome of SARS-CoV-2 will evolve in the initial stage of the pandemic, which may have implications for infection control policy (*9*).

Codon usage of viral genes is subject to selection pressures imposed by the gene translation machinery of hosts (*10, 11*). Spontaneous mutation, fine-tuning of translation kinetics, and evasion of immune recognition together shape the evolutionary trend of codon usage (*12*). Traditional methods based on relative synonymous codon usage (RSCU)(*13, 14*) count the frequencies of codons of virus genes and evaluate the adaptation to the hosts. However, the sparsity of single nucleotide variants (SNVs) in genomes of SARS-CoV-2 precludes the traditional methods, which were designed for comparison between distant species, from extracting the subtle trend. Here, we introduce a method based on fold change of translational efficiency (FCTE), which accumulates the effects of synonymous SNV distributed sparsely in the genome of SARS-CoV-2 and characterize the adaptation to hosts in the epidemic in China.

## RESULTS

### Density of SNV in the coding regions of SARS-CoV-2

In 12 ORFs of SARS-CoV-2, there are on average only 0.021 nonsynonymous sites and 0.012 synonymous sites per 1000 nt per genome (Fig. 1A). The numbers of synonymous and nonsynonymous sites in a coding region are both correlated with the length of the coding region (Fig. 1B). Compare to the average mutation density, ORF1ab:A, which is the coding region of the ORF1ab gene upstream of the −1 frameshift site, contains slightly more synonymous sites, while ORF1ab:B, which is downstrem of the −1 frameshift site, contains less synonymous sites.

**Fig. 1.**
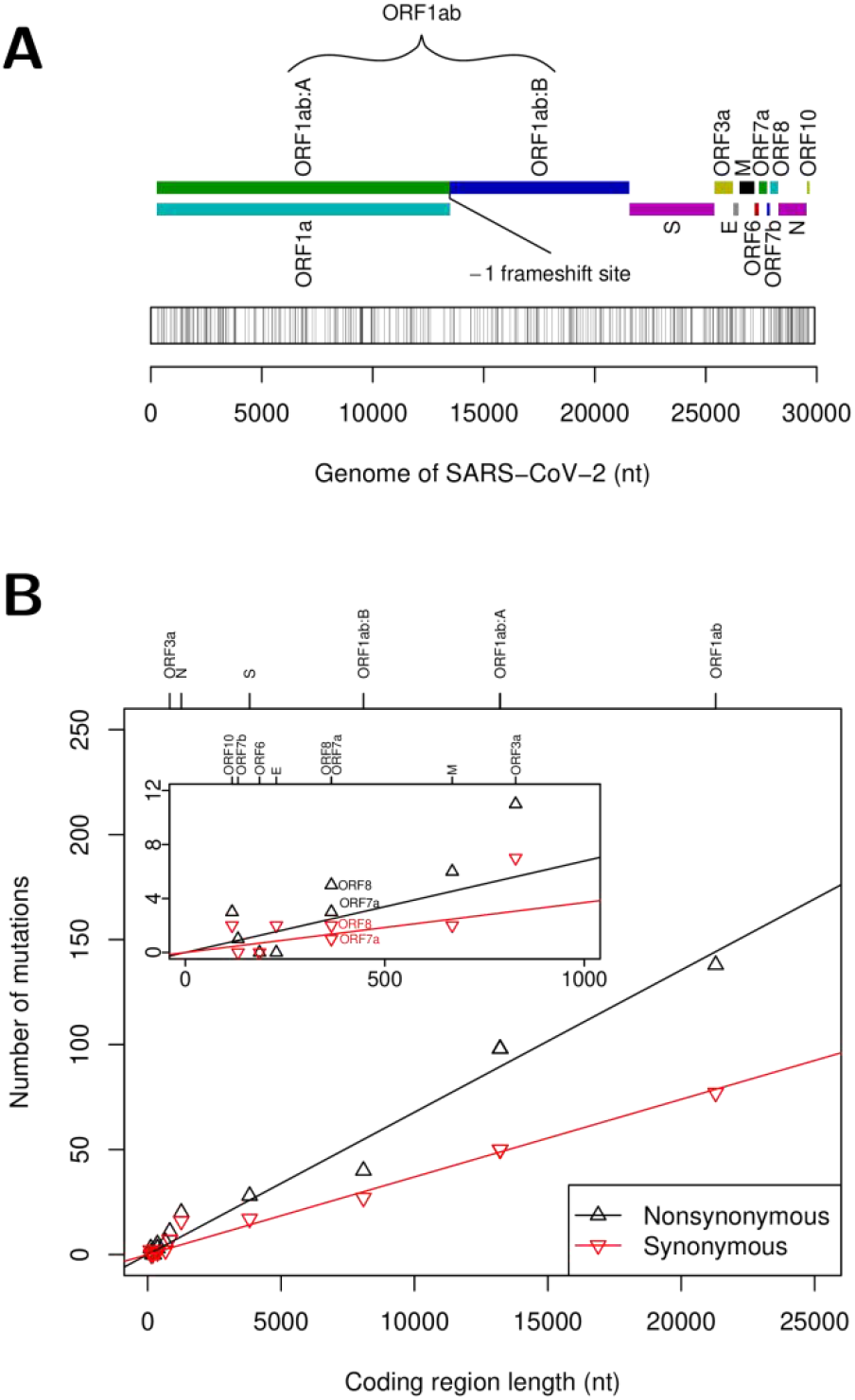
Variant sites in coding regions of the genome of SARS-CoV-2. **(A)** Organization of coding regions in the genome and locations of variant triplet sites. **(B)** Numbers of nonsynonymous and synonymous sites in coding regions.

### FCTE of synonymous mutation

FCTE measures the effect of mutation on the translation of the residing gene. We estimated FCTE by the fold change of codon usage frequencies of the codon pair before and after a synonymous mutation (Table S1). The codon usage frequency was calibrated by the codon frequency averaged over a repertoire of genes weighted by their expression levels in the type II alveolar (AT2) cells of lung tissue, which is probably the target cells of SARS-CoV-2(*15*). Two topologies of phylogenetic trees of the genomes of SARS-CoV-2 were examined. The first topology is star-like, in which the central ancestor is the consensus sequence (Fig. 2A). The second topology is a maximum likelihood tree rooted at the earliest collected genome, in which a mutation is defined as a pair of codon states in the parent and child nodes (Fig. S1, Fig. 2B).

**Fig. 2.**
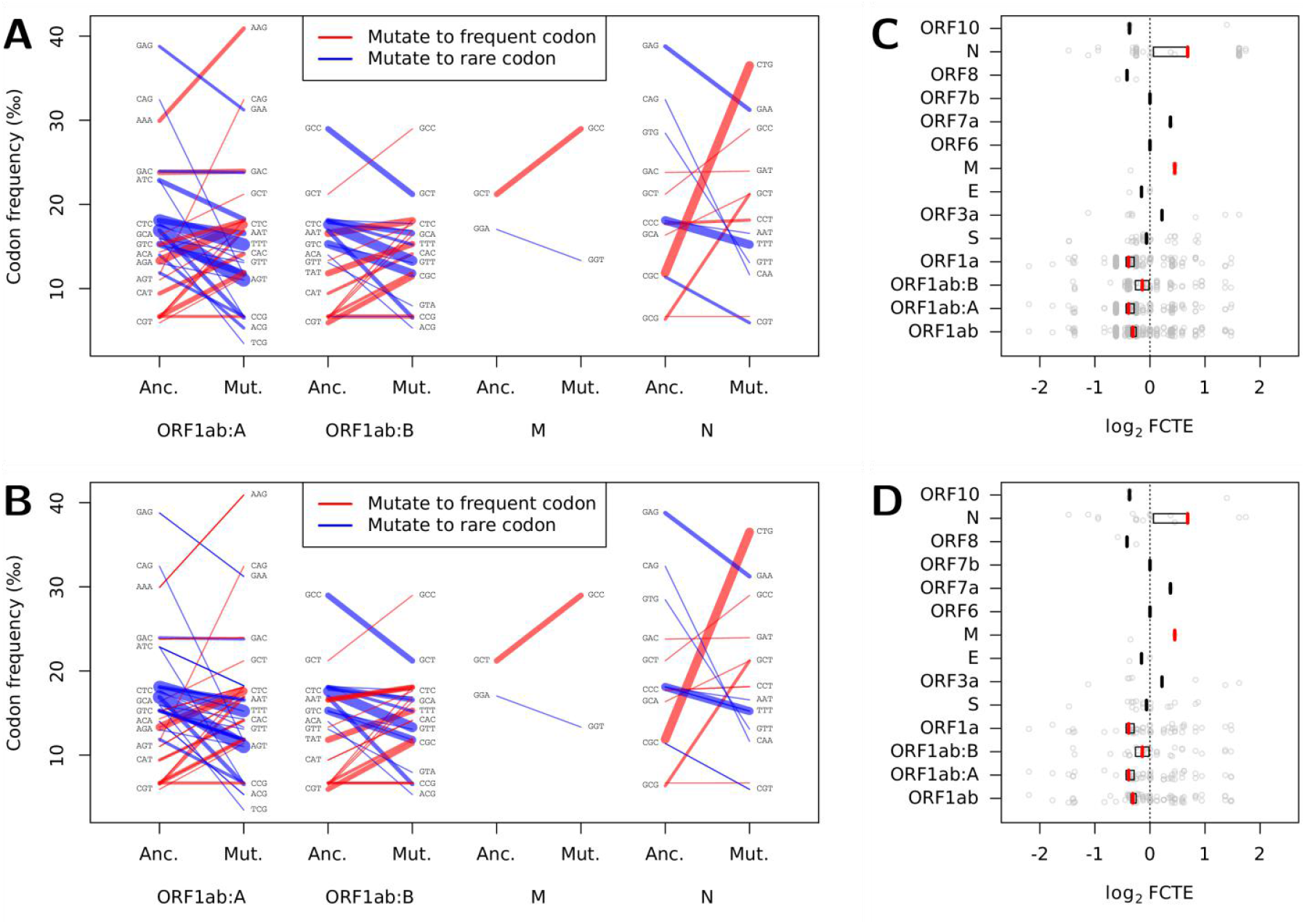
Synonymous sites and fold change of translational efficiency (FCTE). **(A)** Pairs of ancestral (Anc.) versus mutational (Mut.) states of synonymous sites for the star-like tree in the ORF1ab:A and ORF1ab:B fragments and the M and N genes. **(B)** Pairs of ancestral (Anc.) versus mutational (Mut.) states of synonymous sites for the maximum likelihood tree rooted at the earliest collected genome in the ORF1ab:A and ORF1ab:B fragments and the M and N genes. The width of a segment is roughly in proportion to the number of the same mutations in different genomes in the star-like tree or to the prosperity of a mutation in the maximum likelihood tree. Some labels of codons are neglected for clarity. **(C)** FCTE for the star-like tree. **(D)** FCTE for the maximum likelihood tree. Gray points indicate log_2_ FCTE values of mutations, whose pseudomedians are indicated by vertical bars. For coding regions where the Wilcoxon tests reject the null hypothesis that the log_2_ FCTE is zero, the vertical bars are colored red and black boxes are added indicating the nonparametric 95% confidence intervals (except the M gene).

FCTE of mutations were grouped by the residing coding regions and the effects of mutations on the translation were evaluated. The Wilcoxon tests yield almost the same *p* values, pseudomedians, and nonparametric 95% confidence intervals for a coding region considered in different topologies, suggesting that FCTE is insensitive to the topologies of the phylogenetic trees (Fig. 2, C and D). Despite of the ORF6 and ORF7b genes which have no synonymous sites, evolutionary stability of translational efficiency of the S, ORF3a, E, ORF7a, ORF8, and ORF10 genes withstands the Wilcoxon tests. For the M and N genes, the pseudomedians of log_2_ FCTE are 0.45 and 0.68, respectively, suggesting that during the evolution in the epidemic, the population of SARS-CoV-2 benefits from mutations toward codons favored by the hosts, which possibly accelerate the translation of these genes. In the ORF1ab gene, synonymous sites have evolved toward unfavored codons, especially in the ORF1ab:A fragment.

### Evolutionary trends of ORF1ab

To demonstrate the relationship between log_2_ FCTE of mutations and the collection times of the genomes in China, we fitted linear models to mutations in the ORF1ab:A and ORF1ab:B fragments and the full-length ORF1ab gene (Fig. 3). The evolutionary trend of the ORF1ab:A fragment is parallel to the growth of the epidemic (Fig. 3A). The ORF1ab:B fragment shows an evolutionary trend opposite to the ORF1ab:A fragment. A summation of ORF1ab:A and ORF1ab:B, however, obscures the evolutionary trend of the full-length ORF1ab. Examined in context of the maximum likelihood tree rooted at the earliest collected genome, the opposite evolutionary trends of ORF1ab:A and ORF1ab:B are even significant, i.e., the p values of the fitted linear models are both below 0.05 and the correlation coefficients get larger (Fig. 3B).

**Fig. 3.**
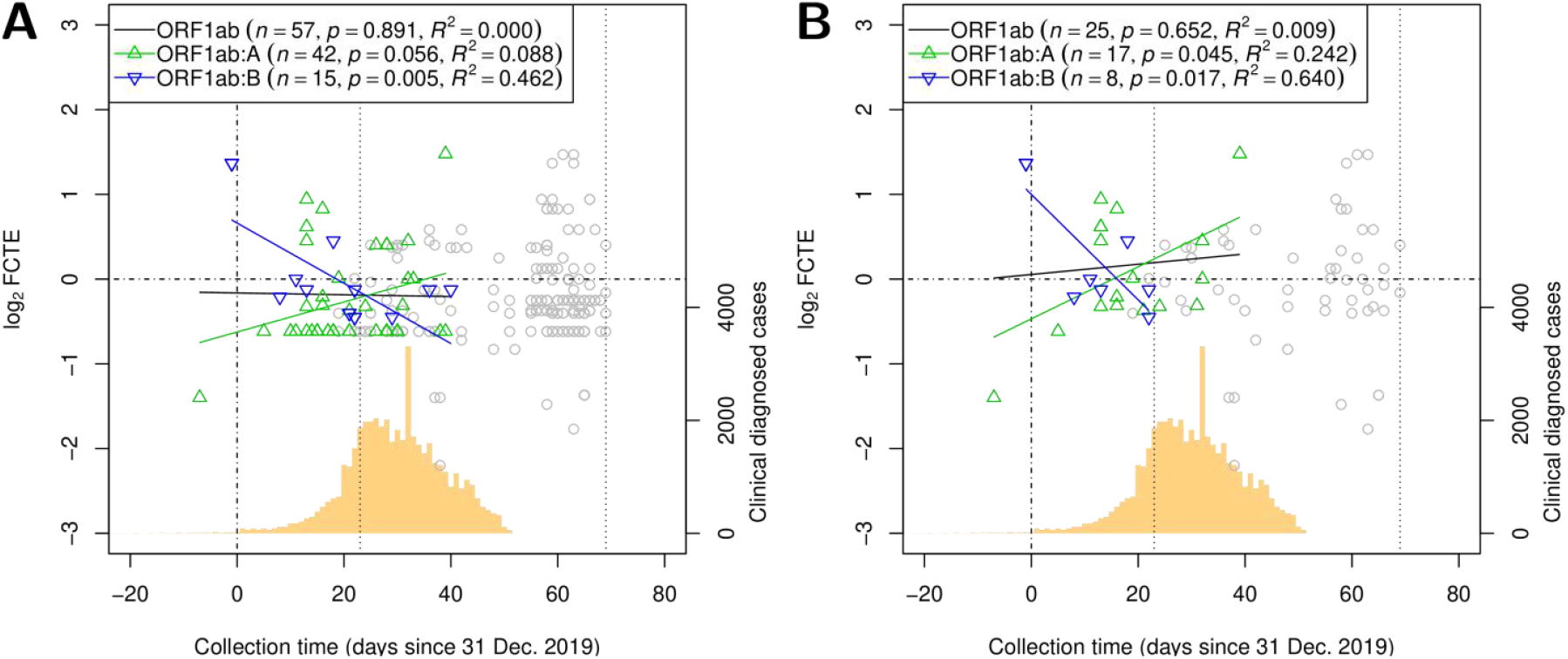
Evolutionary trends of log_2_ fold change of translational efficiency (FCTE) of synonymous sites in the ORF1ab gene. **(A)** log_2_ FCTE for the star-like tree plotted against collection time of the genomes. **(B)** log_2_ FCTE for the maximum likelihood tree plotted against collection time of the genomes. Mutation in genomes collected in China are highlighted with green up-pointing triangles for mutations in ORF1ab:A and blue down-pointing triangles for mutations in ORF1ab:B, with fitted lines. Black lines are fitted to both the green and blue triangle. The numbers of mutations (n), p values of the linear models, and correlation coefficients (R2) are shown. Samples collected elsewhere are shown as gray circles. Vertical dotted lines indicate the dates of Wuhan lockdown (23 Jan. 2020) and Italy lockdown (9 March 2020). The backgrounded histogram shows the numbers of clinical diagnosed cases in Wuhan reported by WHO (*4*).

### Estimation of FCTE for average lung tissue

We estimated FCTE by codon usage frequencies calibrated for average lung tissue (Table S1). The values of codon usage frequencies are similar to those for the AT2 cells. The FCTE of genes (Fig. S2, A, B, E, and F) and the evolutionary trend of the two fragments of ORF1ab (Fig. S3, A and B) are similar to previous results calibrated for the AT2 cells, except that the statistical tests against the evolutionary tendency of the N gene and the evolutionary trend of ORF1ab:A become non-significant (Fig. S2, E and F, and Fig. S3B). We also estimated FCTE by codon usage frequencies calibrated for human genes without weighting (*16*). Although the values of codon usage frequencies are similar to those for the AT2 cells and for average lung tissue, the statistical tests against the evolutionary tendency of the M and N gene and the evolutionary trend of ORF1ab:A are non-significant (Fig. S2, G and H, and Fig. S3D).

## DISCUSSION

Usage of synonymous codons is non-random and non-uniform, the so-called codon bias, the extend of which varies among genes and species (*17*). Codon bias distinguishes expression levels of genes(*13, 18*). Codon usage of the viral genes might be subject to selection pressures imposed by the gene translation machinery of the host, because the virus relies on ribosomes and tRNAs of the host cells(*12*). For human, genes encoding tRNA of various codons are scattered throughout all but the Y chromosome. Relative tRNA abundance, as a result of the number of tRNA genes and the expression levels of these tRNA genes, have been proved correlated significantly to the codon usage of highly expressed genes in a tissue-specific manner(*19*). Since we do not have the tRNA abundance data for lung tissues, we estimate the change of translational efficiency by the change of codon usage in expressed genes in the AT2 cells and in average lung tissue. Although such an estimation may be even more suitable for evaluation of adaptation of virus genome, because the codon usage frequency also reflects the stability of interactions between mRNA and anticodon in addition to tRNA abundance(*12*), an direct measurement of tRNA abundance may help to further address various effects of translation kinetics.

FCTE in the M and N genes confirm the adaptation of the genome of SARS-CoV-2 to the target tissue (*12, 20*). Translation products of the M and N genes, both the most abundant structural proteins, interact with the genomic RNA intimately in the viral replication and virion packaging (*21, 22*). By contrast, synonymous sites in the S gene suggest no tendency of translational efficiency, which can be explained by the fact that the S protein plays no roles in the replication and only a passive role in the packaging. Selection pressure detected in the M and N genes may nullify previous studies on the virus evolution based on the nonsynonymous-to-synonymous site ratio (Ka/Ks test), though speculations on the evolution of the S gene still holds.

The ORF1ab gene contains a ribosomal −1 frameshift site (Fig. 1A)(*23*). When the frameshifting is urged, the product is the ORF1ab protein, which can be further processed to produce RNA-dependent RNA polymerase (RdRp). The newly generated RdRp protein starts synthesizing the complementary strand from the 3’ end of the genomic RNA and disrupts long distance pairing of RNA secondary structure, preventing the frameshifting in later translation(*24*). So the ribosome will encounter a stop codon soon and the product is the ORF1a protein contains no RdRp. By such a mechanism the virus encourages RdRp to replicate the genomic RNA which is free of hindering ribosomes. Although synonymous site may changing the RNA secondary structure(*23*), we observed only one synonymous site appeared in the stem-loop regions overlapped with the ORF1ab gene in one genome(*20*). A tendency of mutation to rare codons shown by the negative value of log_2_ FCTE of the ORF1ab:A fragment (Fig. 2, C and D) imply that SARS-CoV-2 has slowed down the translation and enhanced the −1 frameshifting, producing redundant RdRp and distracting virus replication. According to the SIR (susceptible-infected-recovered) epidemiological model, such a tendency may be evolutionarily beneficial to the virus when the distraction of virus replication assists immune evasion or reduces the mortality rate of the hosts, but is essentially evolutionarily unfavored if there were multiple infections and competition (*25*). In such cases, the genome will be cheated by peers without rare codons in the ORF1ab:A fragment, which produce few RdRp proteins and replicate their genomic RNA with redundant RdRp produced by the cheatee. Parallel to the growth of the epidemic(*4*) which might increase the possibility of multiple infections, the tendency of mutation toward rare codons in the ORF1ab:A region was weakening in the SARS-CoV-2 epidemic in China (Fig. 3B). Hinged by the fine-tuning of FCTE of the ORF1ab:A region, multiple infections break the balance between RdRp production and genome replication, and may drive the evolution of efficient replication. Because high abundance of virus in an infected host increases the virulence and delays the host curing the infection(*25*), prevention against multiple infection seems to be a precondition for taming the virus, may provide a new theoretical basis for social distancing in addition to reducing the infected population (*26*).

## MATERIALS AND METHODS

### Codon usage frequency in the AT2 cells and in lung

The codon usage frequency was calibrated by the averaged frequency of codons in a repertoire of genes weighted by the expression levels. SARS-CoV-2 mainly infects lung tissue, and probably targets the type II alveolar (AT2) cells(*15*). A dataset of single-cell transcriptome of human lung tissue (GEO: GSE122960)(*27*) was re-analysis with R package Seurat version 3.1.4(*28*) following a previous study(*15*). Briefly, high-quality cells from eight donor lung biopsies were extracted, which have 200 to 6,000 detected genes and their mitochondrial gene content was <10%, and genes detected in less than three cells were discarded. The gene-cell matrices from eight donor lung biopsies were integrated by SCT normalization method with 2000 anchor genes. By a shared 20-nearest neighbor graph constructed with the first 30 principle components, the cells were clustered at a resolution of 0.15. The AT2 cluster was identified by comparing the marker genes found by the function FindAllMarkers() to marker genes reported by Reyfman *et al*.(*27*) For each AT2 cell, the frequency of codons in high-quality cells were counted according to coding DNA sequences of GRCh38.84 provided by ENSEMBL(*29*). Codon usage frequency in an AT2 cell was calculated by the frequency of codons averaged among genes weighted by the expression levels. Codon usage frequency in all AT2 cells was averaged and used in subsequent analysis.

In addition, we also calculated codon usage frequency for the average lung tissue. A dataset of the expression level of transcripts measured as TPM in lung tissue was downloaded from GTEx portal (version 8)(*30*). The corresponding transcript sequences were obtained from GENCODE version 26(*31*). Codon usage frequency in a biopsy was calculated by the averaged frequency of codons weighted by the expression levels. Codon usage frequency in lung tissue was averaged over 578 biopsies, of which the autolysis score was 0 or 1, the donor was 20 to 59 years old, and the death Hardy Scale was 1 or 2.

### Multiple sequence alignment and phylogenetic analysis

We downloaded 623 sequences from GISAID’s EpiCoV™ (http://www.gisaid.org, last visit on 13 March 2020, External Data S1), and obtained the sequence and annotations of the reference genome of SARS-CoV-2 from NCBI (accession: NC_045512) (*1*). A multiple sequence alignment (MSA) of 516 genome sequences (sequences of more than 29000 nt) was built using MUSCLE version 3.8.31(*32*) with default setting. The MSA was trimmed, removing nucleotides preceding the first ORF (ORF1ab) and following the last ORF (ORF10). For quality control, we discarded genomes of which the collection date was ambiguous or the sequence contained any gaps or more than one unresolved nucleotide (symbol “N”), which also included genomes collected from pangolins and bats. The refined MSA consists of 317 sequences.

A maximum likelihood phylogenetic tree of the 317 sequences was built using RAxML-NG version 0.9.0(*33*) with GTR+G substitution model and bootstrapping with MRE-based convergence criterion up to 1000 replicates. The tree was essentially unrooted, and we rooted the tree at the earliest collected sample (EPI_ISL_402123) (Fig. S1).

### Synonymous triplet sites

Codon triplets on 12 ORFs and two fragments of ORF1ab, the fragments of ORF1ab:A and ORF1ab:B, totally 14 coding regions were examined (Fig. 1). In the 12 ORFs, 366 SNV triplet sites among 317 genomes distribute in the coding regions uniformly (Fig. 1A). The numbers of nonsynonymous and synonymous sites in a coding region are both correlated with the length of the coding region (Fig. 1B).

In columns of the MSA of a coding region, the frequency of codons was counted. For a synonymous triplet site, the states before and after a mutation were defined by two topologies of the phylogenetic tree of the virus genomes. The first topology was star-like, in which all 317 genomes as nodes were individually connected to the root, which was the consensus sequence of the 317 genomes. A pair of states of a synonymous site before and after the mutation was the consensus codon versus the minority codon, which was the codon in a few virus genomes different from the consensus. Identical minority codons in different genomes were counted separately as pairs of states of the mutation. Minority codons that contained any partial ambiguity symbols were neglected. The star-like topology might blur the evolutionary trend, because it counted descendant of a synonymous mutation several times.

The second topology of the phylogenetic tree of the virus genomes was a maximum likelihood tree rooted at the earliest collected genome (Fig. S1). For a synonymous triplet site, the maximum likelihood ancestor states in internal nodes of the tree were reconstructed using R package phangorn version 2.5.5 (*34*). Pairs of states of a synonymous site before and after the mutation were defined for edges whose two ends had different codons, i.e., the sum of absolute difference of reconstructed probability of states of codons was greater than 1. The ancestral and mutational states of the edge were the codon of the parent and the child nodes, respectively. The number of descendants of the child node indicates the prosperity of the mutation.

### Fold change of translational efficiency (FCTE)

FCTE characterizes the tendency of codon usage bias in coding regions. In this study, we estimated FCTE by the fold change of codon usage frequency before and after a synonymous mutation (Fig. 2, A and B). Synonymous sites of a coding region were grouped by the residing coding regions, and we performed a one-sample exact Wilcoxon test to see if translational efficiency of the coding region was evolutionary stable (Fig. 2, C and D).

We plotted log_2_ FCTE of mutations in the ORF1ab gene against collection times of the virus genomes. For displaying the parallel development of FCTE of mutations with epidemic (Fig. 3), we extracted from World Health Organization’s report on COVID-19 the epidemic curve of clinical diagnosed cases, which include laboratory confirmed cases, in Wuhan (*4*).

In addition to the FCTE estimated by codon usage frequency of AT2 cells, we also calculated the FCTE by codon usage frequency for average lung tissue and for all human genes without weighting (Fig. S2 and S3).

### Statistical Analysis

We implemented scripts for finding synonymous triplet sites and calculating FCTE with R version 3.6.2. To evaluate the evolutionary stability of the translational efficiency of a coding region, synonymous sites of the coding region were grouped by the residing coding regions, and a one-sample exact Wilcoxon test was performed to see if the distribution of the logarithm to base 2 (log_2_) of FCTE of a region was symmetric about zero. Along with the Wilcoxon tests, pseudomedians and the nonparametric 95% confidence intervals were reported except for the M gene, of which the small sample size precluded the estimation of a 95% confidence interval (Fig. 2, C and D). To demonstrate the evolutionary trend of translational efficiency of the ORF1ab gene, the mutations were fitted by linear models. We reported the correlation coefficients (*R^2^*) of the linear models and performed *F*-tests to show the overall significance (Fig. 3).

## General

We acknowledge the researchers originating and submitting the sequences from GISAID’s EpiCoV^™^ on which this study is based (External Data S1).

## Funding

This work was supported by National Key R&D Plan (No. 2018YFC1315400.2, to HL).

## Author contributions

QH and HL conceived the study. QH and HG contributed implementation of the methods and data analysis. LZ, XC, and SH contributed the data collection. QH and HG prepared the manuscript. All authors read and approved the manuscript.

## Competing interests

Authors declare no competing interests.

## SUPPLEMENTARY MATERIALS

**Fig. S1.**
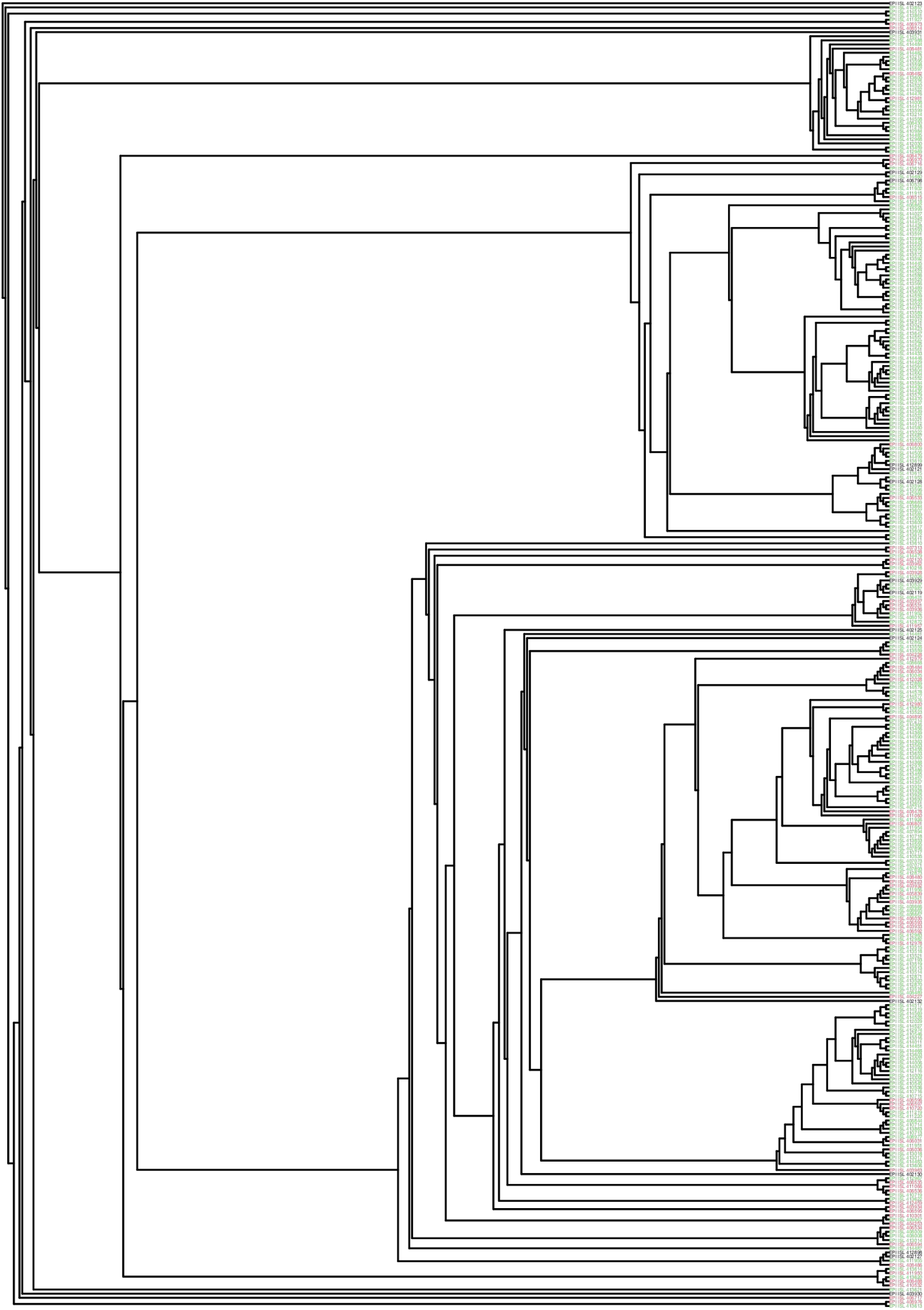
The maximum likelihood tree of 317 virus genomes rooted at the earliest collected virus (EPI_ISL_402123). Accession ID from GISAID’s EpiCoV™ database is colored black if the virus was collected before 31 Dec. 2019, red if between 31 Dec. 2019 and 23 Jan. 2020 (the date of Wuhan lockdown), green if between 23 Jan. 2020 and 9 March 2020 (Italy lockdown), and blue if after 9 March 2020.

**Fig. S2.**
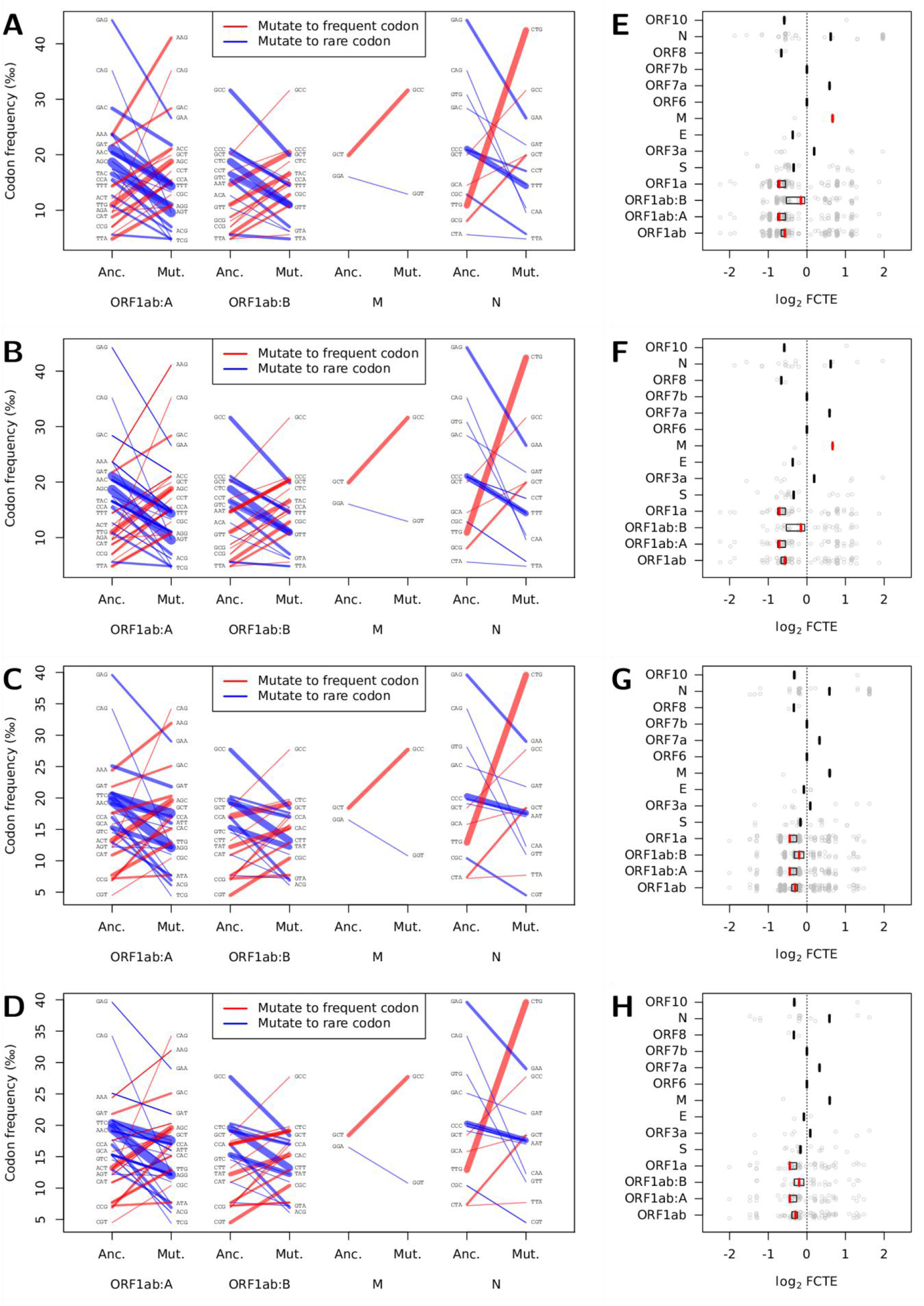
Synonymous sites and fold change of translational efficiency (FCTE) with gene expression levels calibrated for lung tissue and without tissue specificity. **(A)** Pairs of ancestral (Anc.) versus mutational (Mut.) states of synonymous sites for the star-like tree in the ORF1ab:A and ORF1ab:B fragments and the M and N genes with gene expression levels calibrated for lung tissue. **(B)** Pairs of ancestral (Anc.) versus mutational (Mut.) states of synonymous sites for the maximum likelihood tree rooted at the earliest collected genome in the ORF1ab:A and ORF1ab:B fragments and the M and N genes with gene expression levels calibrated for lung tissue. **(C)** Pairs of ancestral (Anc.) versus mutational (Mut.) states of synonymous sites for the star-like tree in the ORF1ab:A and ORF1ab:B fragments and the M and N genes without tissue specificity. **(D)** Pairs of ancestral (Anc.) versus mutational (Mut.) states of synonymous sites for the maximum likelihood tree rooted at the earliest collected genome in the ORF1ab:A and ORF1ab:B fragments and the M and N genes without tissue specificity. The width of a segment is roughly in proportion to the number of the same mutations in different genomes in the star-like tree or to the prosperity of a mutation in the maximum likelihood tree. Some labels of codons are neglected for clarity. **(E)** FCTE for the star-like tree with gene expression levels calibrated for lung tissue. **(F)** FCTE for the maximum likelihood tree with gene expression levels calibrated for lung tissue. **(G)** FCTE for the star-like tree without tissue specificity. **(H)** FCTE for the maximum likelihood tree without tissue specificity. Gray points indicate log_2_ FCTE values of mutations, whose pseudomedians are indicated by vertical bars. For coding regions where the Wilcoxon tests reject the null hypothesis that the log_2_ FCTE is zero, the vertical bars are colored red and black boxes are added indicating the nonparametric 95% confidence intervals (except the M gene).

**Fig. S3.**
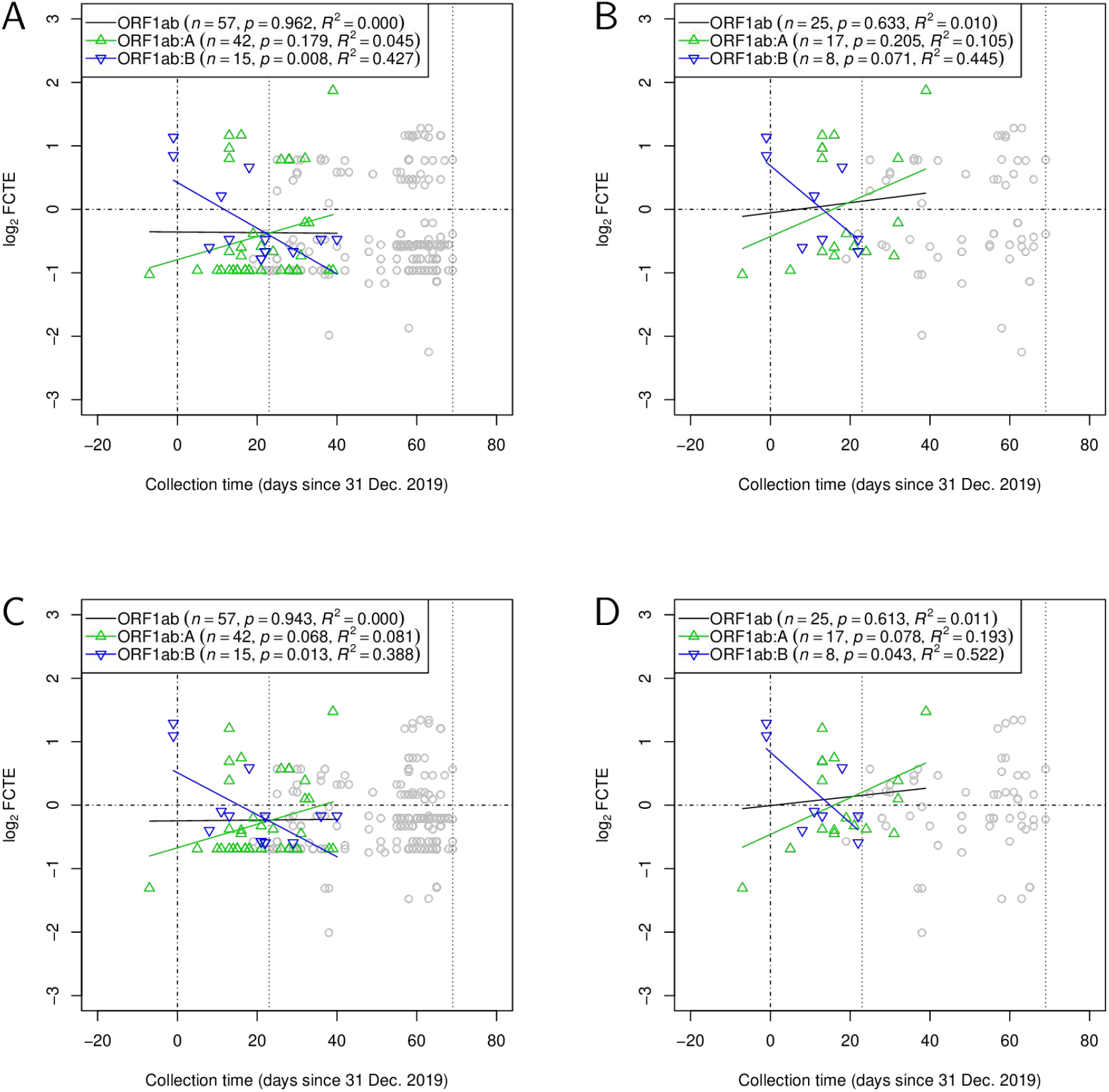
Evolutionary trends of log_2_ fold change of translational efficiency (FCTE) of synonymous sites in the ORF1ab gene with gene expression levels calibrated for lung tissue and without tissue specificity. **(A)** log_2_ FCTE for the star-like tree plotted against collection time of the genomes with gene expression levels calibrated for lung tissue. **(B)** log_2_ FCTE for the maximum likelihood tree plotted against collection time of the genomes with gene expression levels calibrated for lung tissue. **(C)** log_2_ FCTE for the star-like tree plotted against collection time of the genomes without tissue specificity. **(D)** log_2_ FCTE for the maximum likelihood tree plotted against collection time of the genomes without tissue specificity. Mutation in genomes collected in China are highlighted with green up-pointing triangles for mutations in ORF1ab:A and blue down-pointing triangles for mutations in ORF1ab:B, with fitted lines. Black lines are fitted to both the green and blue triangle. The numbers of mutations (*n*), *p* values of the linear models, and correlation coefficients (*R*^2^) are shown. Samples collected elsewhere are shown as gray circles. Vertical dotted lines indicate the dates of Wuhan lockdown (23 Jan. 2020) and Italy lockdown (9 March 2020).

**Table S1.**
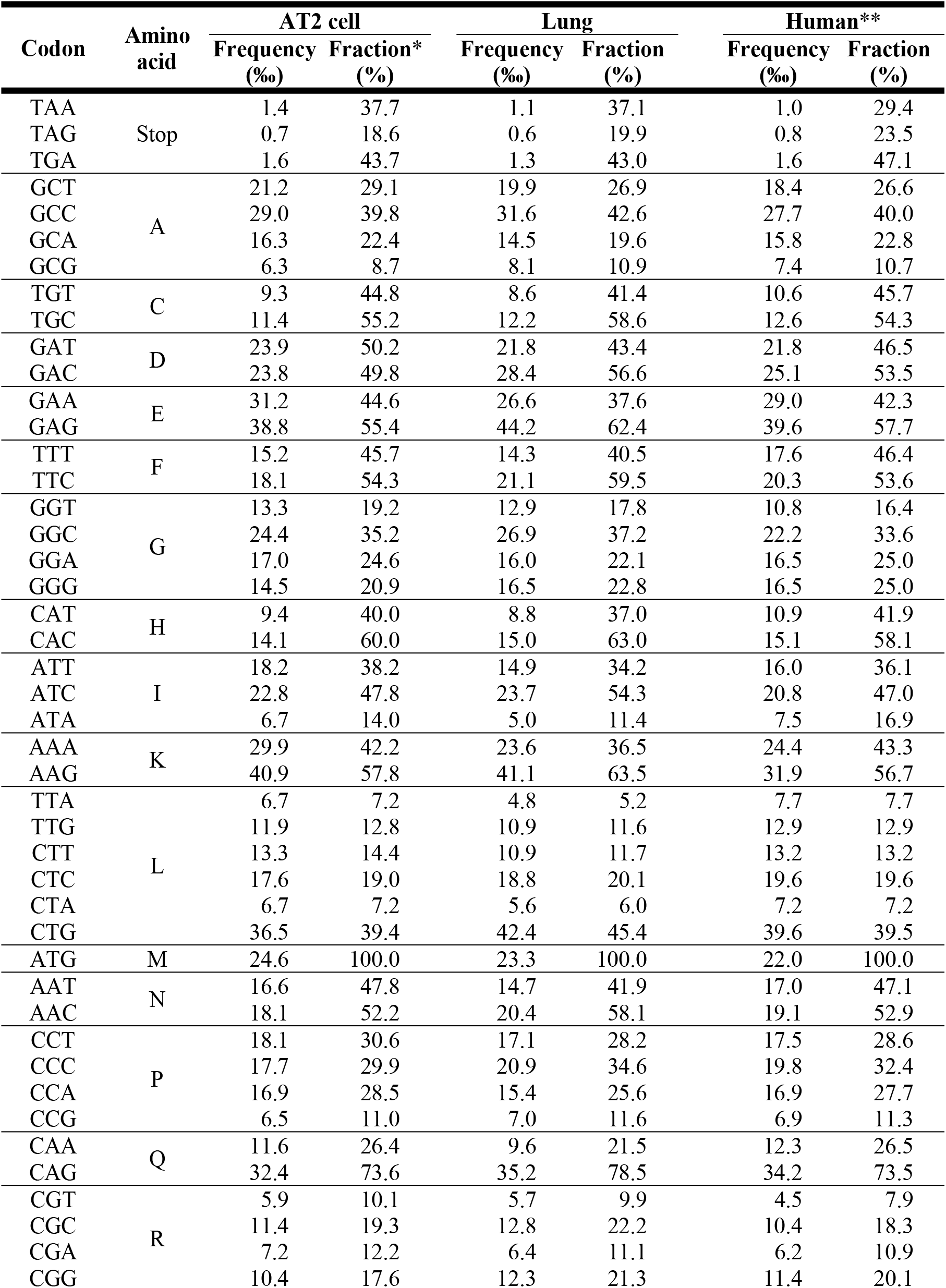
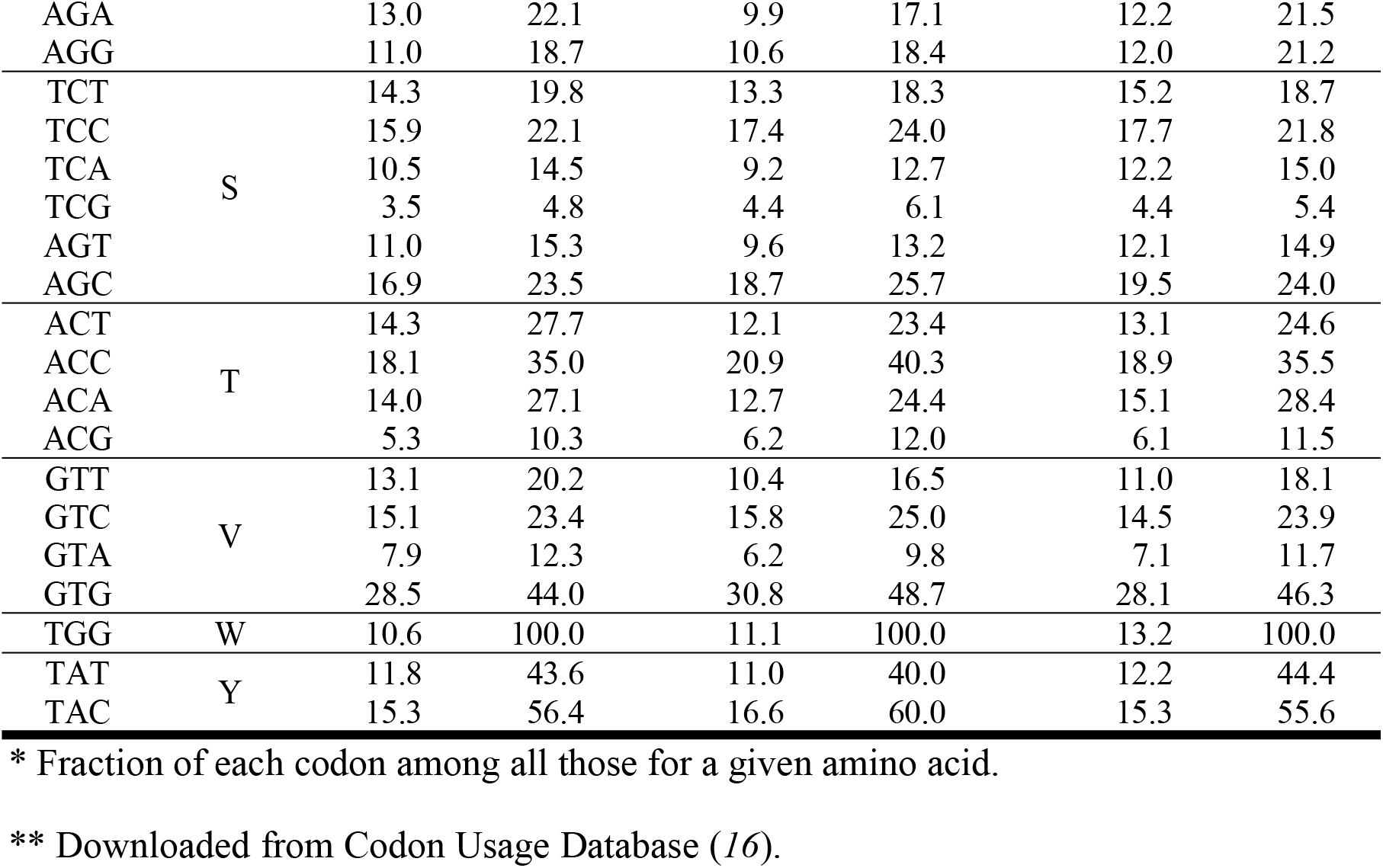
Codon usage frequency table used for estimation of fold change of translational efficiency (FCTE).

**Data S1. Sequences from GISAID’s EpiCoV™ on which this study is based.** The last column “317 genomes” indicates whether a genome was used in constructing the maximum likelihood phylogenetic tree and examining the synonymous triplet sites.

## Notes

### Competing Interest Statement

The authors have declared no competing interest.

